# Generation of an induced pluripotent stem cell line, LCSBi021-A, from a Parkinson’s disease patient harboring the missense mutation c.213C>G (p.Ser71Arg) in the *RAB32* gene

**DOI:** 10.64898/2026.10.20.760406

**Authors:** Ioanna Boumpoureka, Vyron Gorgogietas, Elizabet Petkovski, Rashi Halder, François Massart, NCER-PD Consortium, Rejko Krüger

## Abstract

Dermal fibroblasts were obtained from a 40-year-old male patient with Parkinson’s disease (PD), harboring the c.213C>G (p.Ser71Arg) variant in the *RAB32* gene. Induced pluripotent stem cells (iPSCs) were generated using the CytoTune™-iPS 2.0 Sendai Reprogramming Kit (Thermo Fisher Scientific). The presence of the c.213C>G base exchange in exon 1 of *RAB32* was confirmed by Next Generation Sequencing. The iPSCs are free of genomically integrated reprogramming genes, express pluripotency markers, display *in vitro* differentiation potential into all three germ layers and exhibit karyotypic integrity. Our iPSC line is available for research purposes to dissect molecular mechanisms of neurodegeneration in PD.

## Resource Table

**Resource Table:**
|  |  |
| --- | --- |
| Unique stem cell line identifier | LCSBi021-A |
| Alternative name(s) of stem cell line | N/A |
| Institution | Luxembourg Centre for Systems Biomedicine |
| Contact information of the distributor | Prof. Dr. Rejko Krüger |
| Type of cell line | iPSCs |
| Origin | Human |
| Additional human source info required for human ESC or iPSC | Age at the biopsy: 40 years old<br>Age of disease onset: 27 years old<br>Sex: male<br>Ethnicity: Caucasian |
| Cell Source | Dermal fibroblasts |
| Clonality | Clonal |
| Method of reprogramming | Transgene-free<br>CytoTune™-iPS 2.0 Sendai Reprogramming Kit (Thermo Fisher Scientific) |
| Genetic Feature(s) | c.213C>G (p.Ser71Arg) in the *RAB32* gene |
| Evidence of the reprogramming transgene loss (including genomic copy if applicable) | Transgene-free, confirmed by PCR |
| Associated disease | Parkinson's disease (OMIM #168600) |
| Gene/locus | Ras-related protein Rab-32 (*RAB32*) /<br>GRCh38:CM000668.2 |
| Date archived/stock date | March 2026 |
| Cell line repository/bank | Luxembourg Centre for Systems Biomedicine |
| Ethical approval | Ethical approval for the development of and research pertaining to patient-derived cell lines have been given by informed consent for the academic research project (CNER #201411/05): “Disease modeling of Parkinson's disease using patient-derived fibroblasts and induced pluripotent stem cells” (DiMo-PD). |

## Resource utility

Parkinson’s disease (PD) is the fastest-growing neurodegenerative disorder worldwide (Bloem et al., 2021). RAB32 p.S71R was recently identified to cause autosomal dominant PD in multiple individuals worldwide (Gustavsson et al., 2024; Hop et al., 2024). However, the exact mechanisms underlying RAB32-mediated neurodegeneration remain unclear. Our patient-derived iPSC line will serve as a tool to study RAB32 pathophysiology *in vitro*.

## Resource Details

Dermal fibroblasts were obtained from a 40-year-old male PD patient, harboring the point mutation p.S71R in RAB32 (c.213C>G base exchange in exon 1 of the *RAB32* gene). The patient exhibited right-sided tremor and postural instability at the age of 27 and received a PD diagnosis at the age of 31. He is currently presenting with rigidity, bradykinesia, postural tremor, hyposmia, micrographia, and anxiety, but is not receiving dopaminergic therapy. Fibroblasts at passage 3 (p3) were reprogrammed using the CytoTune™-iPS 2.0 Sendai Reprogramming Kit, according to the manufacturer’s instructions. Four weeks after transduction, iPSC colonies started emerging, which were picked manually and transferred to separate Geltrex-coated plates, to preserve clonality. One clone was further expanded to establish the LCSBi021-A line, which showed typical iPSC morphology, creating compact colonies with distinct borders and well-defined edges (**Fig. 1A**). PCR confirmed the loss of reprogramming factors at p9 (**Fig. 1B**), and Next Generation Sequencing (NGS) confirmed the presence of the mutation in both fibroblasts and generated iPSCs (**Fig. 1G**). Pluripotency was validated with immunocytochemistry (ICC) for SSEA-4 and SOX2 (**Fig. 1D**), and RT-qPCR for *NANOG, DNMT3B*, and *OCT4* (**Fig. 1E**). Normal molecular karyotype was confirmed by a CGH microarray (**Fig. 1C**). Moreover, the iPSCs are capable of directed *in vitro* differentiation into the three germ layers, as shown by ICC for the markers OTX2 (ectoderm), SOX17 (endoderm), and Brachyury (mesoderm) (**Fig. 1F**). An STR marker analysis confirmed the identity of the line (submitted to the journal’s archive). Lastly, both fibroblasts and the iPSC line tested negative for mycoplasma contamination (**Fig. S1A**) (Table 1).

**Table 1:** Characterization and validation.

| Classification | Test | Result | Data |
| --- | --- | --- | --- |
| **Morphology** | Brightfield photography | Compact colonies with sharp edges | Fig. 1A |
| **Characterization of undifferentiated state** | Marker expression analysis | Robust nuclear staining of SOX2 and cytoplasmic staining of SSEA-4, as seen by ICC | Fig. 1D |
|  |  | Positive expression of the pluripotency markers *OCT4*, *NANOG*, and *DNMT3B*, as seen by RT-qPCR | Fig. 1E |
| **Genomic stability** | CGH microarray | No chromosomal aneuploidies or large structural aberrations. The small copy-number variants detected (14 intervals, 23 kb–2.6 Mb) largely map to regions of common copy-number polymorphism or segmental duplication. | Fig. 1C |
| **mtDNA analysis** | N/A | N/A | N/A |
| **Cell line identity verification** | STR analysis | Extremely strong support of identity between patient-derived fibroblasts and generated iPSCs | Submitted to the journal's archive |
| **Mutation analysis** | Next Generation Sequencing | c.213C>G mutation present in *RAB32* exon 1 | Fig. 1G |
|  | Sanger sequencing |  | Fig. S1B |
| **Microbiology and virology** | Mycoplasma test | Negative | Fig. S1A |
| **Characterization of the differentiated state** | Directed *in vitro* tri-lineage differentiation | Expression of markers representative of ectodermal (OTX2), endodermal (SOX17), and mesodermal (Brachyury) formation, as seen by ICC. | Fig. 1F |
| **Donor screening** | HIV 1 + 2 Hepatitis B, Hepatitis C | Not performed | N/A |
| **Genotype additional info** | Blood group genotyping | Not performed | N/A |
|  | HLA tissue typing | Not performed | N/A |

## Materials and Methods

### Cell culture and reprogramming

Primary fibroblasts were cultured in DMEM supplemented with 10% FBS and 1% penicillin/streptomycin. Reprogramming was performed at passage 3 using the CytoTune-iPS 2.0 Sendai Reprogramming Kit (Thermo Fisher Scientific, Cat# A16517), following the fibroblast reprogramming feeder-free protocol, with the following multiplicity of infection (MOI): KOS MOI = 5, hc-Myc MOI = 5, hKlf4 MOI = 3. One week after transduction, cells were replated on Geltrex (Thermo Fisher Scientific, Cat# A1413302)-coated 6-well plates and cultured in mTeSR™ Plus (StemCell Technologies, Cat# 100–0276). Undifferentiated iPSC colonies were manually picked into Geltrex-coated 24-well plates and then expanded into larger well formats. Passaging was done using 0.5 mM EDTA in PBS. Fibroblasts and iPSCs were cultured at 37°C, 5% CO_2_, under humidified atmosphere.

### PCR for loss of reprogramming factors

Total RNA was purified from cells at p9 using the RNeasy Mini Kit (Qiagen, Cat# 74104), according to the manufacturer’s instructions. Transcriptor High Fidelity cDNA Synthesis Kit (Roche, Cat# 5081955001) was used to synthesize cDNA. To assess the loss of reprogramming vector, cDNA was amplified by PCR with the DreamTaq PCR Master Mix (Thermo Fisher, Cat# K1081) using primers specific for the Sendai reprogramming vectors (Table 2). cDNA from fibroblasts lysed four days after transduction were used as a positive control. The amplification products were resolved in a 1% agarose gel, as seen in Fig. 1B.

**Table 2:**
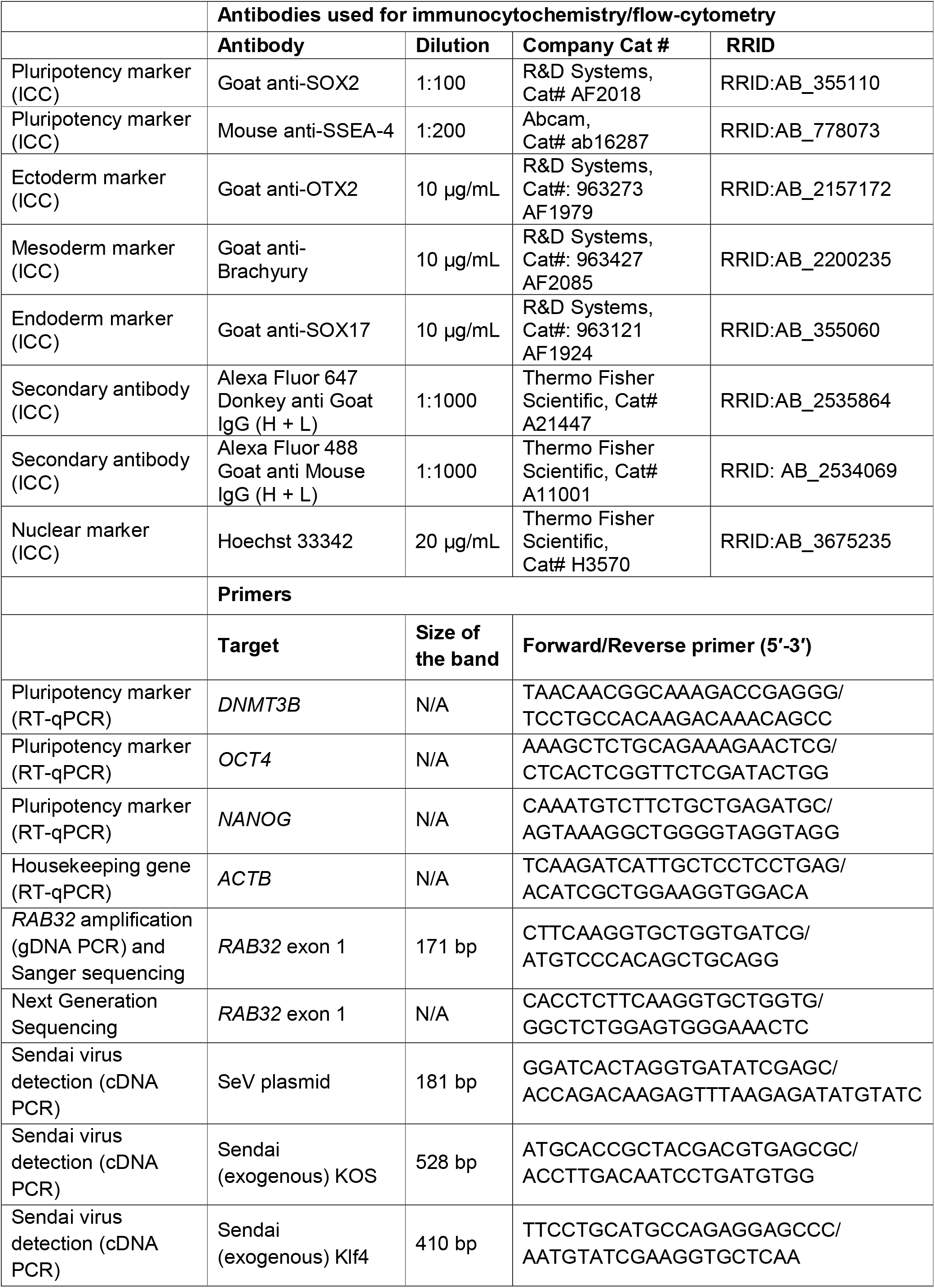
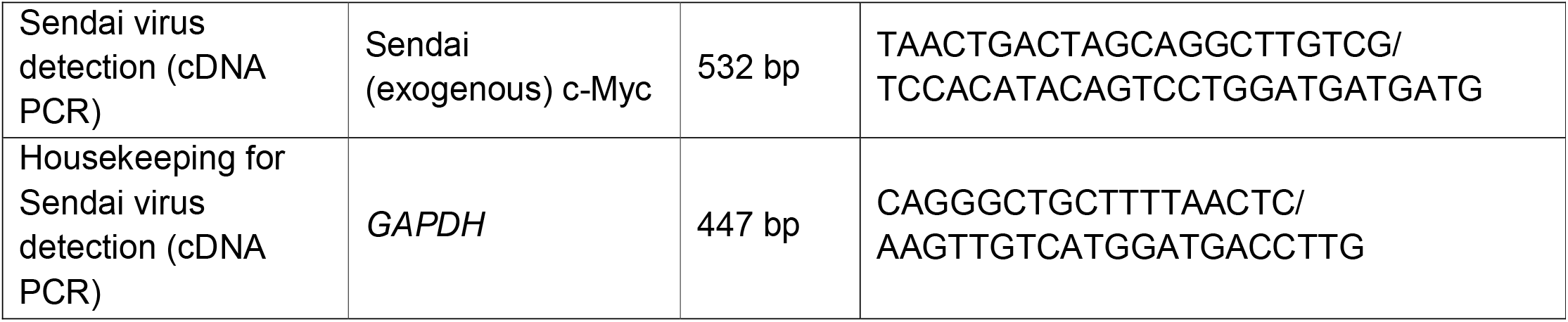
Reagent details.

### Next Generation Sequencing

gDNA was isolated from iPSCs and fibroblasts, using the DNeasy Blood & Tissue Kit (Qiagen, Cat# 69504) and submitted to LCSB’s Genomics Platform (RRID:SCR_021931) for targeted NGS. A region of *RAB32* exon 1 encompassing c.213 was PCR-amplified and sequenced as 2 × 151 bp paired-end reads on an Illumina iSeq 100, yielding 6,297 (fibroblasts) and 7,958 (iPSCs) read pairs. Reads were compared to the *RAB32* reference sequence (NM_006834), and the base at c.213 was called in read pairs in which the forward and reverse reads agreed at that position. The c.213C>G variant was detected in both fibroblasts (p5) and iPSCs (p6), as presented in Fig. 1G.

### Sanger sequencing

*RAB32* exon 1 was amplified from gDNA using the primers listed in Table 2 and Sanger-sequenced at Microsynth. Owing to the high GC content of exon 1, only partial forward-strand reads were obtained, as presented in Fig. S1B.

### Pluripotency evaluation with ICC

iPSCs at p6 were fixed with 4% paraformaldehyde in PBS for 20 min at RT, followed by permeabilization and blocking with 0.1% Triton X-100 (Sigma, Cat# X100) and 2% bovine serum albumin (BSA; Roth, Cat# 8076.4) in PBS for 1 h at RT. Subsequently, cells were incubated overnight at 4°C with primary antibodies against SOX2 and SSEA-4 in PBS with 2% BSA, and the next day they were washed three times with PBS and incubated with secondary antibodies and 20 μg/mL of the nuclear marker Hoechst 33342 (Thermo Fisher Scientific, Cat# H3570) for 2 h at RT (Table 2). Images were acquired with the Zeiss Axio Observer 7 microscope (Carl Zeiss MicroImaging GmbH) and are presented in Fig. 1D, with a scale bar of 100 μm.

### Pluripotency evaluation with RT-qPCR

At p6, total RNA extraction followed by cDNA synthesis were performed as mentioned above. Quantification by qPCR was performed using the PowerTrack™ SYBR Green Master Mix for qPCR (Thermo Fisher Scientific, Cat# A46012), and primers against *DNMT3B, OCT4*, and *NANOG* (Table 2). *ACTB* was used as a housekeeping gene. Samples were run on the LightCycler (Roche) following the standard cycling mode suggested by the manufacturer (1 cycle of enzyme activation in 95°C for 2 min and 40 cycles of denaturation in 95°C for 15sec and annealing/extension in 60°C for 60sec). cDNA from the patient’s fibroblasts was used as a negative control, and cDNA from LCSBi016-A was used as a positive control (Yoldi Bergua et al., 2026). Data in Fig. 1E are shown as fold change normalized to LCSBi016-A iPSCs.

### Chromosomal analysis

iPSCs at p7 were karyotyped with a comparative genomic hybridization (CGH) microarray at Laboratoire National de Santé (LNS). DNA was isolated with the DNeasy Blood & Tissue Kit (QIAGEN, Cat# 69504) and treated with 100 μg/mL RNase A. iPSC and control DNA (Agilent Human Male Reference DNA) were then prepared for aCGH using the SureTag Complete DNA Labeling protocol (v8.0, Agilent). DNA was then comparatively hybridized to a SurePrint G3 Human CGH Microarray 180 K (Agilent). After washing and scanning, the array data were analyzed and visualized using Cytogenomics (Agilent), as presented in Fig. 1C.

### Directed tri-lineage *in vitro* differentiation

The ability of iPSCs to differentiate *in vitro* into the three germ layers was tested at p7 with the Human Pluripotent Stem Cell Identification Kit (R&D Systems, Cat# SC027B), according to the manufacturer’s instructions. Immunocytochemistry confirmed the expression of OTX2 (ectoderm), Brachyury (mesoderm), and SOX17 (endoderm) (Table 2). Images are presented in Fig. 1F, with a scale bar of 100 μm.

### Mycoplasma test

After 72 h of cell culture, the medium was collected and tested for mycoplasma contamination using the PlasmoTest™ Mycoplasma Detection Kit (Invivogen, Cat# rep-pt1), according to the manufacturer’s instructions. Testing was performed regularly. The experiment presented in Fig. S1A is from p8 (iPSCs) and p6 (fibroblasts).

### Identity analysis

The STR reference genetic profiles of the fibroblasts and iPSCs were characterized using the PowerPlex® Fusion 6C (Promega) PCR amplification system at LNS. Fragment analysis was done on an ABI3500 (Life Technologies), and the resulting data were analyzed with GeneMapper ID-X software (Applied Biosystems; RRID:SCR_014290). iPSCs from p4 and fibroblasts from p5 were used for this experiment. Data are submitted to the journal’s archive.

## CRediT authorship contribution statement

**Ioanna Boumpoureka:** Data curation, Formal analysis, Investigation, Visualization, Writing – original draft. **Vyron Gorgogietas:** Investigation, Writing – review & editing. **Elizabet Petkovski:** Investigation, Writing – review & editing. **Rashi Halder**: Investigation, Writing – review & editing. **François Massart:** Investigation, Writing – review & editing. **Rejko Krüger:** Conceptualization, Methodology, Project administration, Supervision, Writing – review & editing.

## Acknowledgements

**Ioanna Boumpoureka** is supported by the Luxembourg Fonds National de Recherche (FNR) within the framework of the NextImmune2 Doctoral Training Unit (PRIDE21/16749720/NEXTIMMUNE2). **Rejko Krüger** is receiving research grants from the Luxembourg Fonds National de Recherche (FNR ID 19007959), the Michael J. Fox Foundation for Parkinson□s Research, the Innovative Medicines Initiative (<u>IMI ID 101034344</u> — EPND) of the European Union, and the European Union’s Horizon Europe research and innovation programs (INTER/PerMed22/17045084/DEEPEN-iRBD).

We would also like to thank all participants of the Luxembourg Parkinson’s Study for their important and continued support to our research. We acknowledge the Luxembourg National Research Fund (FNR/NCER13/BM/11264123) for funding the National Centre of Excellence in Research on Parkinson’s Disease (NCER-PD) until 2023 and the sustained support by the institutional partners of the NCER-PD consortium, the Luxembourg Institute of Health (LIH) as coordinator, the Luxembourg Centre for Systems Biomedicine (LCSB)/University of Luxembourg, the Centre Hospitalier de Luxembourg (CHL), and the Laboratoire National de Santé (LNS). Furthermore, we acknowledge the joint effort of all NCER-PD Consortium members from our partner institutions generally contributing to the Luxembourg Parkinson’s Study.

---

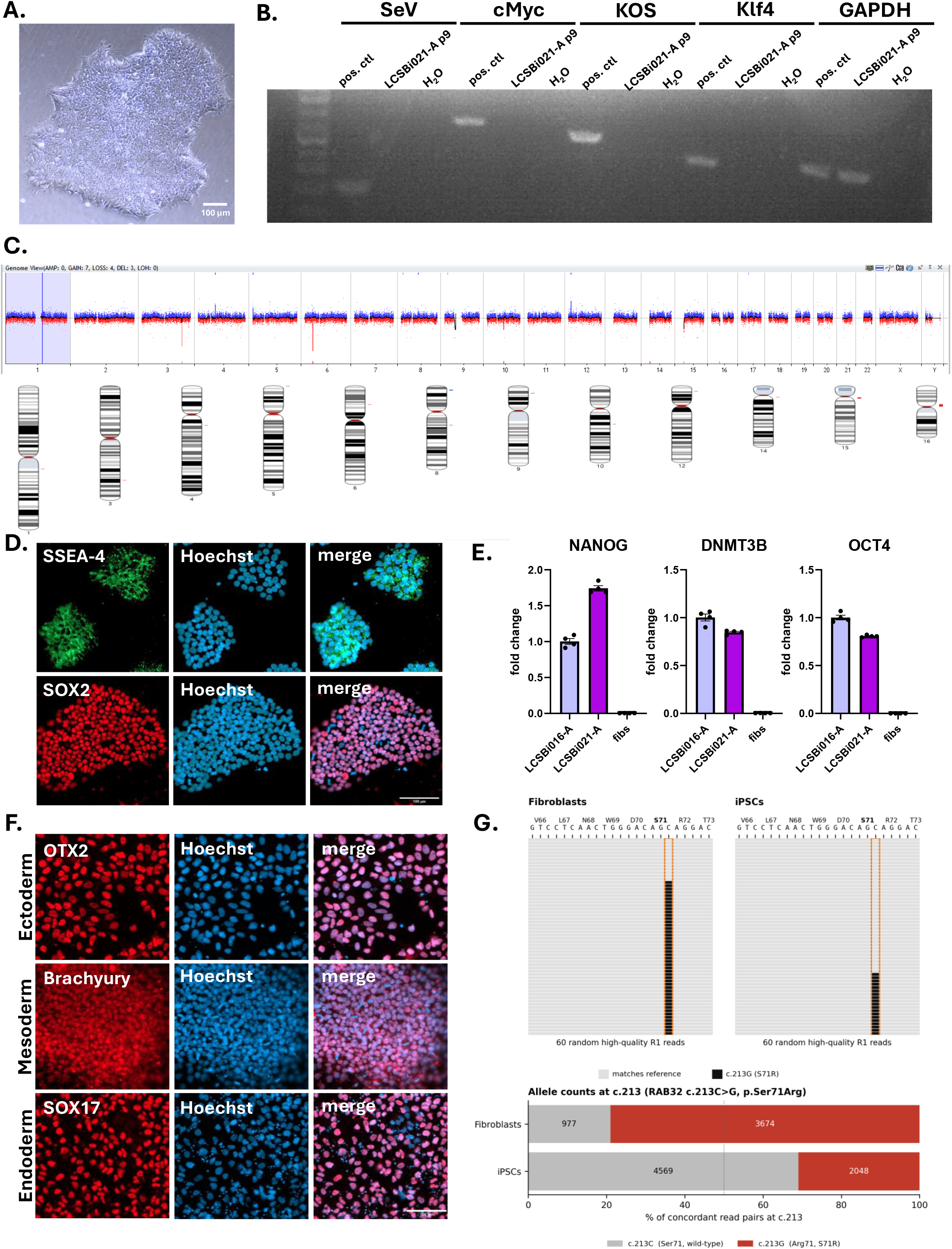

