## Supplementary Figure S1 for "Generation of an induced pluripotent stem cell line, LCSBi021-A, from a Parkinson’s disease patient harboring the missense mutation c.213C>G (p.Ser71Arg) in the *RAB32* gene"

**A.**

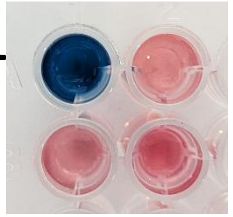


**Samples:**
- **+** positive control
- **-** negative control
- **1** LCSBi021-A iPSCs – passage 8
- **2** Rab32 – fibroblasts – passage 6

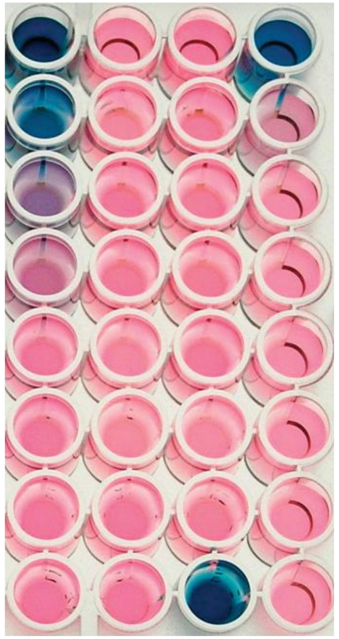

### Reading and Interpretation from the protocol

**Vizualisation of Mycoplasma contamination using PlasmoTest™**

Column 1: Positive and negative controls
Column 2: Uninfected cell line supernatants and M. hyorhinis infected 293PR
Column 3: Uninfected cell line supernatants
Column 4: Dilutions of 293PR supernatant infected with M. hyorhinis

**B.**

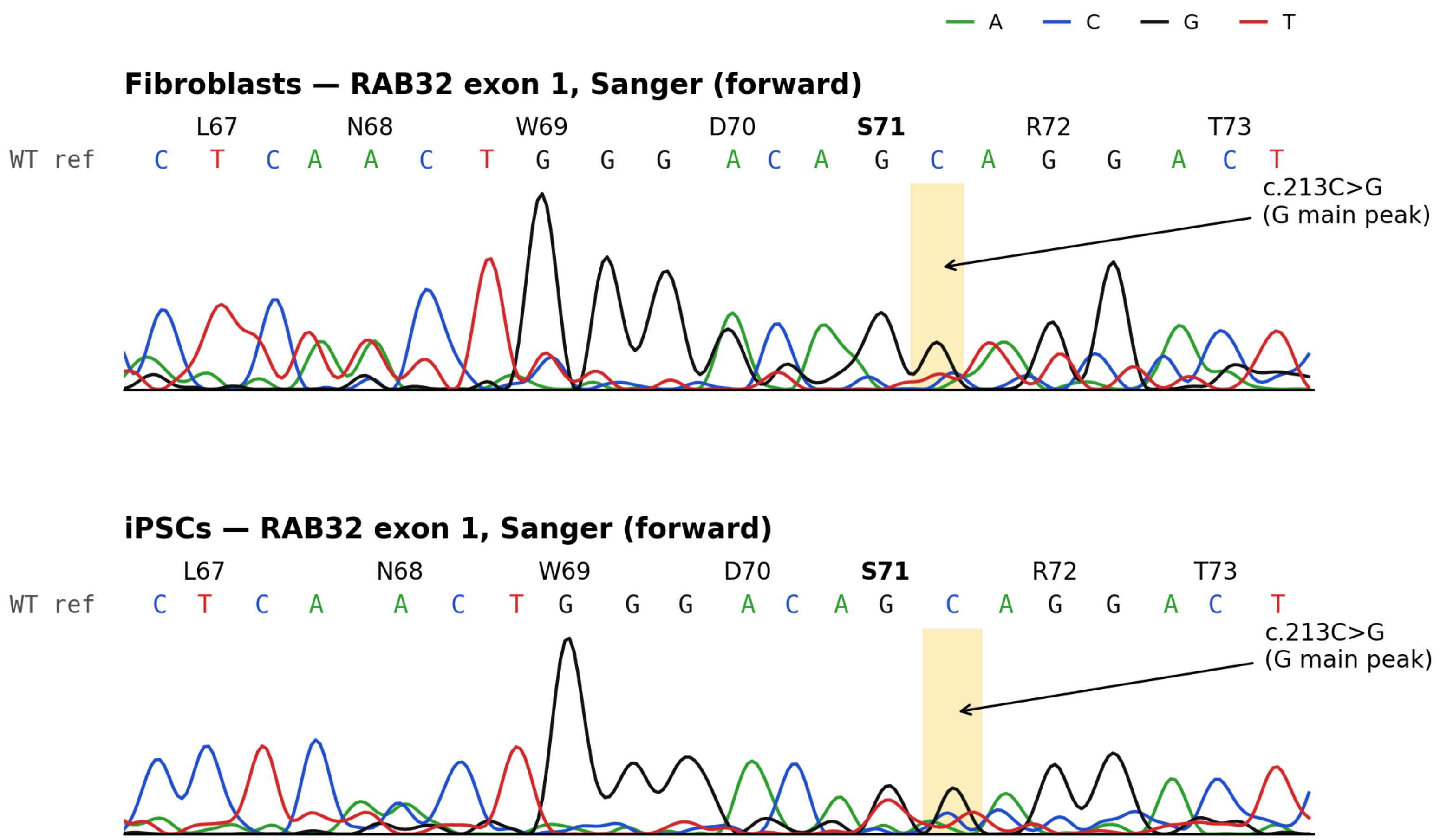


Note on Sanger sequencing: RAB32 exon 1 lies within a GC-rich region (~64% GC), which hindered PCR amplification and Sanger sequencing despite repeated attempts. As a result, trace quality declines around codon 71 and no interpretable reverse-strand reads were obtained. The traces are therefore shown as supporting data only. The presence of c.213C>G in both fibroblasts and iPSCs was established by targeted NGS (Fig. 1G).
